# Yeast *FIT2* homolog is necessary to maintain cellular proteostasis by regulating lipid homeostasis

**DOI:** 10.1101/2020.04.06.027847

**Authors:** Peter Shyu, Wei Sheng Yap, Maria Laura Gaspar, Stephen A. Jesch, Charlie Marvalim, William A. Prinz, Susan A. Henry, Guillaume Thibault

**Author notes:** To whom correspondence should be addressed: Guillaume Thibault: School of Biological Sciences Nanyang Technological University, Singapore, 637551.

## Abstract

Lipid droplets (LDs) have long been regarded as inert cytoplasmic organelles with the primary function of housing excess intracellular lipids. More recently, LDs have been strongly implicated in conditions of lipid and protein dysregulation. The fat storage inducing transmembrane (FIT) family of proteins comprises of evolutionarily conserved endoplasmic reticulum (ER)-resident proteins that have been reported to induce LD formation. Here, we establish a model system to study the role of *S. cerevisiae FIT* homologues (*ScFIT*), *SCS3* and *YFT2*, in proteostasis and stress response pathways. While LD biogenesis and basal ER stress-induced unfolded protein response (UPR) remain unaltered in *ScFIT* mutants, *SCS3* was found to be essential for proper stress-induced UPR activation and for viability in the absence of the sole yeast UPR transducer *IRE1*. Devoid of a functional UPR, *scs3*Δ mutants exhibited accumulation of triacylglycerol within the ER along with aberrant LD morphology, suggesting a UPR-dependent compensatory mechanism for LD maturation. Additionally, *SCS3* was necessary to maintain phospholipid homeostasis. Strikingly, the absence of the *ScFIT* proteins results in the downregulation of the closely-related Heat Shock Response (HSR) pathway. In line with this observation, global protein ubiquitination and the turnover of both ER and cytoplasmic misfolded proteins is impaired in *ScFIT*Δ cells, while a screen for interacting partners of Scs3 identifies components of the proteostatic machinery as putative targets. Taken together, these suggest that ScFIT proteins may modulate proteostasis and stress response pathways with lipid metabolism at the interface between the two cellular processes.

---

Lipid droplets (LDs) have long been regarded as inert cytoplasmic organelles with the primary function of housing excess intracellular lipids. LDs arise from the endoplasmic reticulum (ER) and contain a core of non-polar lipids triacylglycerol (TG) and steryl ester (SE) surrounded by a phospholipid monolayer, with phosphatidylcholine (PC) as the major component (1). More recently, LDs have been strongly implicated in conditions of lipid and protein dysregulation. These conditions are major contributors to the pathophysiology of metabolic diseases and concomitantly activate cellular stress response pathways, namely the unfolded protein response (UPR) and heat shock response (HSR). Increased LD biogenesis has been extensively observed in cells under stress conditions. Introduction of oxidative stressors or the attenuation of antioxidant capacities of cells result in the formation of LDs (2-4). While the extent of UPR activation and the resulting transcriptional profile differs between proteotoxic and lipid stress, a similar increase in lipogenic markers and concomitant LD formation is observed (5-10). It remains to be determined whether LDs contribute to stress induction or if this is reflective of the adaptive role of LDs to mitigate the otherwise deleterious effects of stress. However, all these undoubtedly highlight the complex integration of LDs in stress response pathways. In addition to this, the UPR regulates metabolic pathways to a certain extent under normal physiological conditions (11), and similarly orchestrates the complex transcriptional metabolic reprogramming under ER stress induction (7).

The fat storage-inducing transmembrane (FIT) family of proteins constitutes a group of evolutionarily conserved proteins eponymously named for their role in lipid metabolism and LD formation (12). In mammals, FIT proteins exhibit differential expression patterns, with FIT1 being expressed primarily in cardiac and skeletal muscle tissue, and FIT2 being more ubiquitously expressed. FIT proteins (FIT1/FIT2) are ER-resident proteins with a total of six transmembrane domains and with both N- and C-termini facing the cytosol (12,13). *S. cerevisiae FIT2* homologs (*ScFIT*), *SCS3* and *YFT2*, are predicted to share the same membrane topology. The pioneer study on the FIT proteins has identified its profound effect on the formation and accumulation of LDs both *in vitro* and *in vivo* (12). Transient overexpression of *FIT2* was sufficient to drive the formation of LDs, a process that was later hypothesized to be mediated by the capacity of FIT2 to directly bind and partition TG from the ER into storage in LDs *in vitro* (13). Transgenic expression of the mammalian *FIT2* gene in the budding yeast *S. cerevisiae* (14) and in the plant models *Arabidopsis thaliana* and *Nicotiana tabacum* (15) induced the formation of cytoplasmic LDs. Similarly, transient expression of the *ScFIT* genes in mammalian cells *in vitro* led to the increased formation of LDs (14). A gain-of-function FIT2 mutant was identified with a 3-amino acid mutation within its fourth transmembrane domain, which interestingly houses the most highly conserved amino acid residues from yeast to humans (16). Conversely, it was found that the deletion of *FIT2* greatly compromised LD formation in the model organisms, *Danio rerio* and *Caenorhabditis elegans*, as well as in the pathogenic yeast *Candida parapsilosis* (12,17,18). Similar findings were observed in FIT2-ablated human cells cultured *in vitro* (19). All these reports further reinforced the initial hypothesis of the FIT proteins directly functioning in LD biogenesis.

In contrast, *S. cerevisiae* deletion mutants for either or both *ScFIT* genes retain their capacity to form LDs, with size and number comparable to WT (14,19). Upon closer investigation, it was found that LDs fail to completely bud off from the ER in the absence of the ScFIT proteins (17), presumably through alterations in ER membrane lipid properties (20). These findings have since then led to the investigation of alternative functions of the FIT class of proteins. *SCS3* was initially reported to have a putative role in the regulation of phospholipids (21), a function that was largely unexplored until a large-scale genetic screen reported on the strong interactions between the *ScFITs* and genes involved in phospholipid biosynthesis, including *DGK1* and *PSD1* (14). In-depth analysis of the ScFIT protein sequences have since then revealed the presence of the catalytic site of a lipid phosphatase (22). This finding was further corroborated by a separate study that reported the same observation on the mammalian FIT2 sequence (19). *In vitro* analyses have identified the capacity of mammalian FIT2 to hydrolyze phosphates from phosphatidic acid (PA) and lyso-PA to yield diacylglycerol (DAG) and monoacylglycerol (MAG), respectively. On the other hand, substrates for the lipid phosphatase activity of the ScFIT proteins have yet to be determined. This conserved catalytic function has been associated with the aberrant ER whorling phenotype observed in cells devoid of the FIT proteins (19,22).

Conjointly, several different perspectives now exist on the function of the ScFIT proteins. However, definitive evidence for the function of FIT proteins either in binding and partitioning NLs or influencing NL and phospholipid metabolism in the complex *in vivo* environment remains scarce. Moreover, how any of these functions are relevant in the context of cellular function and broader physiological processes has mostly been left uncharacterized.

In this study, we investigated the role of *SCS3*, identified as one of the downstream UPR target genes. As the UPR transducer Ire1 is essential for viability in the absence of *SCS3*, we generated temperature sensitive allele *scs3-1* to reveal its role without the masking effect of the UPR program. We demonstrated that dysfunctional *SCS3* leads to the accumulation of TG at the ER, to a shift in phospholipids distribution, and the biogenesis of aberrant LD morphology. Furthermore, we identified the interactome of ScFITs of which Scs3 interacts with components of the proteostatic machinery. Next, we demonstrated that *ScFIT* mutants impaired the clearance of ER-associated degradation (ERAD) client proteins, which was exacerbated by lipid imbalance. Together, our data support a model where ScFITs modulate proteostasis and stress response pathways with lipid metabolism at the interface between the two cellular processes.

## Results

### Scs3 is essential for viability in the absence of UPR transducer Ire1

Synthetic genetic array (SGA) analyses revealed the synthetic lethality between *SCS3* and *IRE1*, which encodes for the sole UPR transducer in yeast (14). Additionally, the mutant *scs3*Δ strain has been reported to activate the UPR (23), while *SCS3* was conversely shown to be transcriptionally upregulated upon UPR activation resulting from either proteotoxic stress or lipid bilayer stress (LBS) (7,10,24). As previously reported (17,19,22), Scs3 and Yft2 proteins localize to the ER (Fig. S1A). Herewith, we sought to further understand the role of *SCS3* within the UPR program. First, we monitored the UPR activation using the UPR element (UPRE)-LacZ reporter assay (25). Unexpectedly, no significant UPR activation was observed in *scs3*Δ mutants, contradicting a previous report (23) while being consistent with other findings (14) (Fig. 1A). Similarly, there was no significant UPR activation in *yft2*Δ nor in *ScFIT*Δ. All mutant strains are fit to mount an UPR response upon treatment with ER stress-inducing agent tunicamycin (Tm) although the level of activation was significant lower in *scs3*Δ and *ScFIT*Δ. Similarly, the heat shock response (HSR), a cytosolic proteotoxic stress compensatory pathway, was dampened in *ScFIT*Δ upon heat stress (Fig. S1B and S1C). To complement the UPR assay, we asked if *SCS3* and *YFT2* genes are upregulated in a UPR dependent manner. *SCS3* was significantly upregulated while *YFT2* was mildly upregulated in WT cells treated with Tm (Fig. 1B). As depleting the media of inositol induces the UPR by LBS (10,26,27), we measured the mRNA levels of both genes upon inositol depletion (- ino) and only *SCS3* exhibited a mild but significant upregulation. To further assess the role of *SCS3* during ER stress, we carried out a growth assay. The spotting assay revealed that the lack of *SCS3* exhibits growth defect in the presence of Tm which can be rescued with the overexpression (OE) of *SCS3* (Fig. 1C). Together, these results demonstrate that *SCS3* is essential during ER stress conditions.

**Figure 1.**
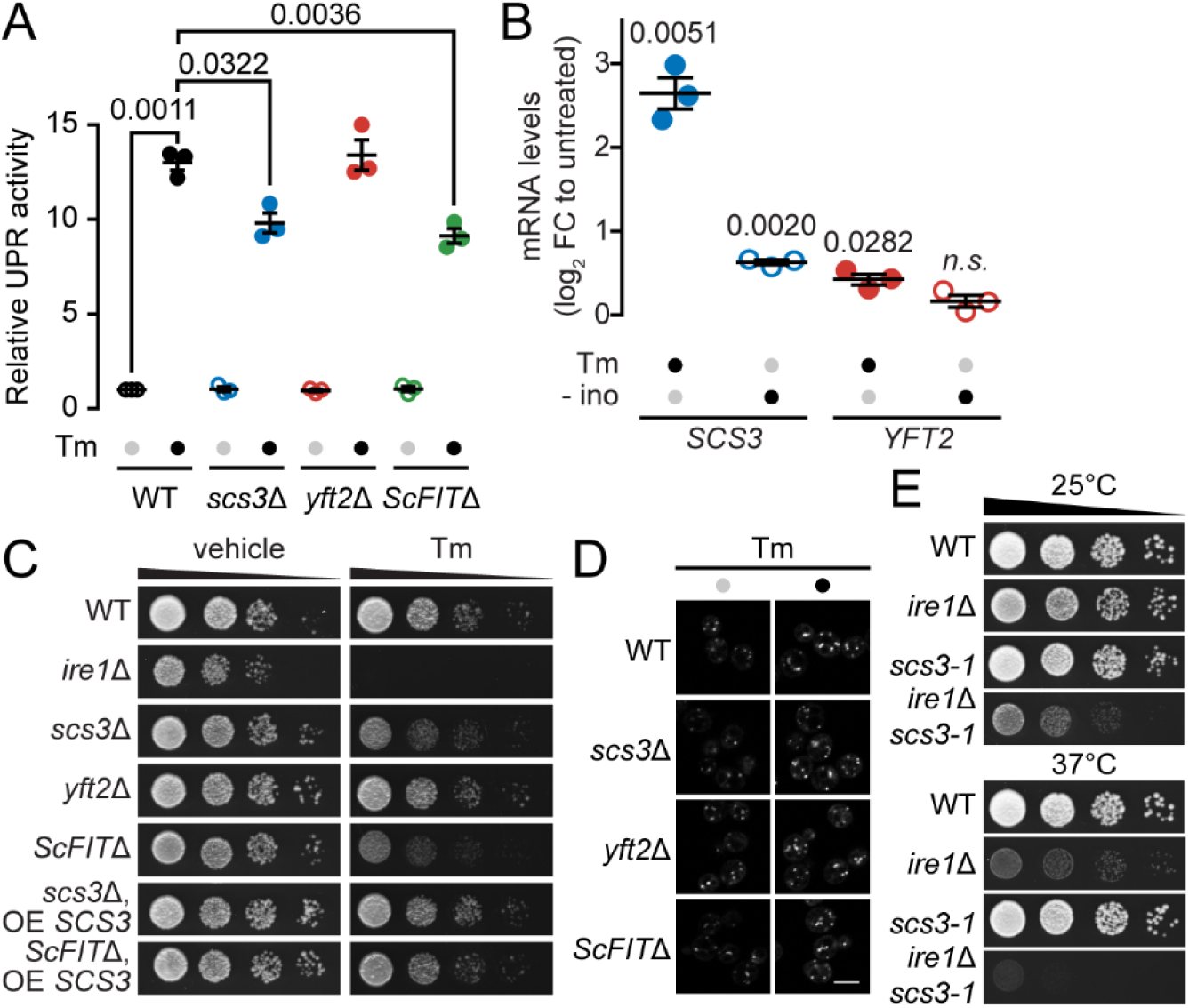
Scs3 is essential for viability in the absence of UPR transducer Ire1. (A) Activation of the UPR was measured by a reporter assay utilizing the expression of the LacZ enzyme under the *CYC1* promoter with the UPR element. Cells were either treated with the *N*-linked glycosylation inhibitor tunicamycin (Tm) or the carrier DMSO for 1 h prior to harvesting for the assay. (B) qPCR results comparing the mRNA levels of *SCS3* and *YFT2* if WT cells with 0.25 μg/ml tunicamycin (Tm) or the absence of inositol (- ino), when indicated. (C) ScFIT mutant strains were grown in selective medium prior to dilution and spotting on vehicle or Tm-supplemented media. Culture plates were incubated at 30°C until colonies are observed. OE, overexpressed. (D) WT and *scs3*Δ mutant cells were treated with Tm for 1 h before LD staining with the fluorescent BODIPY 493/503 dye. Scale bar, 2 µm. Images shown are representative of three independent experiments. (E) Strains in the PER genetic background were grown at 25°C in minimal media from which dilutions were prepared and spotted onto plates. Plates were incubated at the specified temperatures until colonies were observed. Data shown is the mean ± SEM from three independent experiments. Statistical analysis was subjected to paired two-tailed Student’s t-test. *n.s.*, non-significant.

As lipogenic pathways constitute one of the major effectors of the UPR (28) and that LDs are associated with stress conditions, the putative function of Scs3 in LD formation may then provide a rationale for its UPR-dependent transcriptional upregulation. We further hypothesized that the inability of *scs3*Δ mutants to mount a maximal UPR under proteotoxic stress conditions may translate to the impairment of LD formation as part of the stress response. We made use of fluorescent BODIPY 493/503 to stain LDs in *scs3*Δ mutants challenged with Tm-induced stress to evaluate gross changes in LD formation. The mutant cells are still able to form LDs to the same extent as that of WT cells under unstressed and ER stressed conditions (Fig. 1D and S2). Next, we sought to determine the extent by which the UPR compensates for the absence of *SCS3*, particularly in respect to LD formation. However, a *scs3*Δ*ire1*Δ mutant is synthetically lethal, thus rendering conventional deletion strategies unusable in titrating the phenotypic effects of the UPR upon the loss of *SCS3*. To address this, we generated a conditional temperature sensitive *scs3* allele (*scs3-1*) that is functional at a permissive temperature of 25°C but not at the restrictive temperature of 37°C in the absence of *IRE1* (Fig. 1E and S3), using a screening strategy that we previously reported (7,29,30). As *IRE1* is not essential in the absence of *YFT2*, a temperature sensitive allele of *yft2* was not generated (Fig. S3B). This provides strong support for the interdependence between *IRE1* and *SCS3* and that one is required for viability in the absence of the other, consistent with SGA results showing synthetic lethality between these two genes (14).

### Scs3 is essential to maintain lipid homeostasis at the ER and LD morphology

In *scs3*Δ and *yft2*Δ mutants, LDs remain irreversibly tethered to the ER and are wrapped by a membrane. Jacquier *et al*. suggest that, in yeast, LDs always remain connected to the ER (31). We hypothesized that while these knockout strains are fully capable of forming LDs (Fig. 1D), it is possible that these LDs remain tightly associated with the ER, resulting in the disruption of ER lipid homeostasis, as previously reported (17,20). Temperature sensitive allele strains were grown to early log-phase at 25°C followed by a temperature shift of 2h at 37°C. To assess the levels of neutral lipids in the ER, we extracted total lipids from microsomes of the cells and triacylglycerols (TG) was separated by thin layer chromatography (TLC) and quantified by gas chromatography with flame ionization detector (GC-FID) (Fig. 2A). A significant increase in TGs within the microsomal fractions of *scs3-1* strain was only observed in the absence of *IRE1* at the restrictive temperature of 37°C, thereby suggesting that the presence of the UPR exerts a suppressive effect for this phenotype. As LDs were wrapped with ER membranes in *scs3*Δ and *yft2*Δ yeast mutants using inducible TG synthesis system (17), we asked if the UPR plays a role in the budding and morphology of LDs. Transmission electron microscopy (TEM) was performed on *scs3-1* and *ire1*Δ*scs3-1* cells following temperature shift. Strikingly, we observed irregular LD morphology in *scs3-1* and *ire1*Δ*scs3-1* cells (Fig. 2B and S4). The abnormally elongated LDs are embedded in ER. Additionally, LDs were smaller in *ire1*Δ*scs3-1* cells compared to *scs3-1* cells at 37°C. Taken together, our findings suggest that Scs3 is necessary to maintain neutral lipid homeostasis at the ER and that the UPR plays an important role to regulate TAG levels.

**Figure 2.**
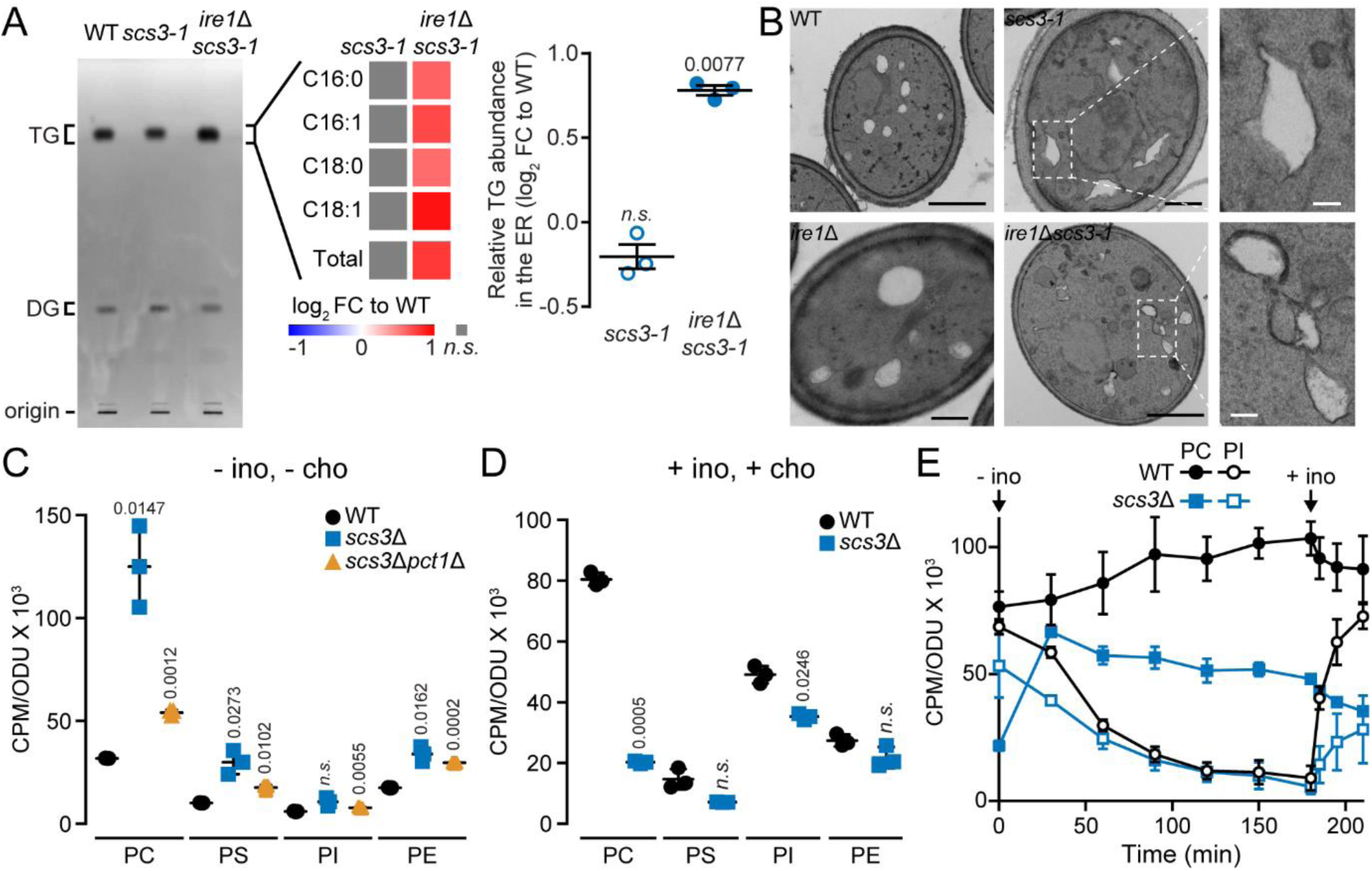
Scs3 is necessary to maintain lipid homeostasis and LD morphology. (A) Strains were grown to log phase at 25°C in minimal media, after which the temperature was shifted to 37°C for 2 h. Extracted microsomal lipids were separated on TLC plates. TG, triacylglycerol; DG diacylglycerol. Changes in TG acyl chain profiles. All FA species were normalised against WT levels. Relative TG abundance was determined by FAME analysis through GC-FID and normalised against WT. (B) Strains were grown to log phase at 25°C in minimal media and shifted to 37°C for 2 h before imaging. Scale bar, 500 nm (zoomed-in images 100 nm). (C) Phospholipid levels of cells grown in media without inositol and choline (- ino, - cho) in the presence of [^32^P]-orthophosphate. Phosphatidylcholine (PC), phosphatidylserine (PS), phosphatidylinositol (PI), and phosphatidylethanolamine (PE) were separated by thin layer chromatography (TLC). (D) Phospholipid levels of cells grown in medial with inositol and choline (+ ino, +cho) treated as in (C). (E) PC and PI levels of cells transferred to media depleted of inositol (- ino) over the course of 3h followed by a reintroduction of inositol (+ ino) for a period of 0.5h. Data shown is the mean ± SEM from three independent experiments. Statistical analysis was subjected to paired two-tailed Student’s t-test. *n.s.*, non-significant.

The transcription of a subset of genes encoding phospholipid biosynthesis enzymes are inhibited in the presence of phospholipid precursors inositol and choline (32-34). Additionally, the absence of inositol in yeast growth media induces the UPR (10,26,27). Therefore, we asked if Scs3 plays a role modulating phospholipid homeostasis in the absence of inositol and choline. We performed lipid analysis of cells grown in media containing inositol and choline to mid-logarithmic phase before being shifted to a 3h incubation in media lacking inositol and choline. Unexpectedly, *scs3*Δ mutant cells contained 4 times the phosphatidylcholine (PC) levels of WT cells (Fig. 2C). PC levels of *scs3*Δ cells were significantly decreased in *pct1*Δ background mutant suggesting that PC is mostly synthesized through the Kennedy pathway in the absence of *SCS3*. Pct1 is a cholinephosphate cytidylyltransferase enzyme. Phosphatidylserine (PS) and phosphatidylethanolamine (PE) were also found to be two times more abundant in *scs3*Δ mutant compared to WT cells while there was no significant difference in phosphatidylinositol (PI) levels. Next, we tested if supplementing the media with inositol and choline will restore phospholipid levels in *scs3*Δ mutant. In contrast to the absence of both lipid precursors, we observed a decrease of about 4 times of PC in *scs3*Δ mutant compared to WT cells (Fig. 2D). There was also a significant decrease of PI in *scs3*Δ mutant. To further understand the role of Scs3 in modulating phospholipids, we measured the levels of PC and PI over the course of 3h of cells grown in media depleted of inositol followed by a 0.5h recovery (+ ino) period. In the presence of choline, *scs3*Δ failed to increase the synthesis of PI upon the re-introduction of inositol while PI level increased rapidly in WT (Fig. 2E). On the other hand, there was a constant decrease of PC levels in *scs3*Δ while PC levels continually increased in WT during the 3h of inositol depletion. Together, these data reveal that Scs3 is essential to maintain phospholipid homeostasis. We speculate that the decrease of PI in *scs3*Δ cells might alter the composition of complex sphingolipids (35) which in turn would induce the UPR (36).

### Scs3 interacts with components of the proteostatic machinery

To gain further insight into the physiological relevance of ScFIT proteins within the cell, we employed the split-ubiquitin based membrane yeast two hybrid (MYTH) screen (37). The reporter moiety was fused to the N-terminal or the C-terminal cytosolic domains of both Scs3 and Yft2. The four bait constructs were validated, and screening conditions for each were optimised with 3’-amino-1,2,4,-triazole (3’-AT) supplementation in the selection media to reduce the occurrence of false positives. Following this, 1,344 colonies were collectively screened for all reporter strains. From these, 664 colonies were positive for bait-prey interaction as manifested by blue colony growth on 5-bromo-4-chloro-3-indolyl-β-D-galactopyranoside (X-gal) supplemented selective media and are designated as putative interactors (Fig. S5). These were further validated for specificity towards the bait protein of interest in comparison to the single-pass human cluster of differentiation 4 (CD4) receptor protein, which served as a negative control. From these, 189 showed specific interactions with the ScFIT proteins. Following sequence analysis, 88 unique protein interactors were identified. Considering the overlap in protein interactors, the MYTH screen identified a total of 73 genuine and unique interactors for the ScFIT proteins, which were further categorized according to cellular functions (Fig. 3). Moreover, our screen results show that more than half of the identified Yft2 protein interactors are shared with that of Scs3, thereby supporting a certain degree of functional redundancy between the two.

**Figure 3.**
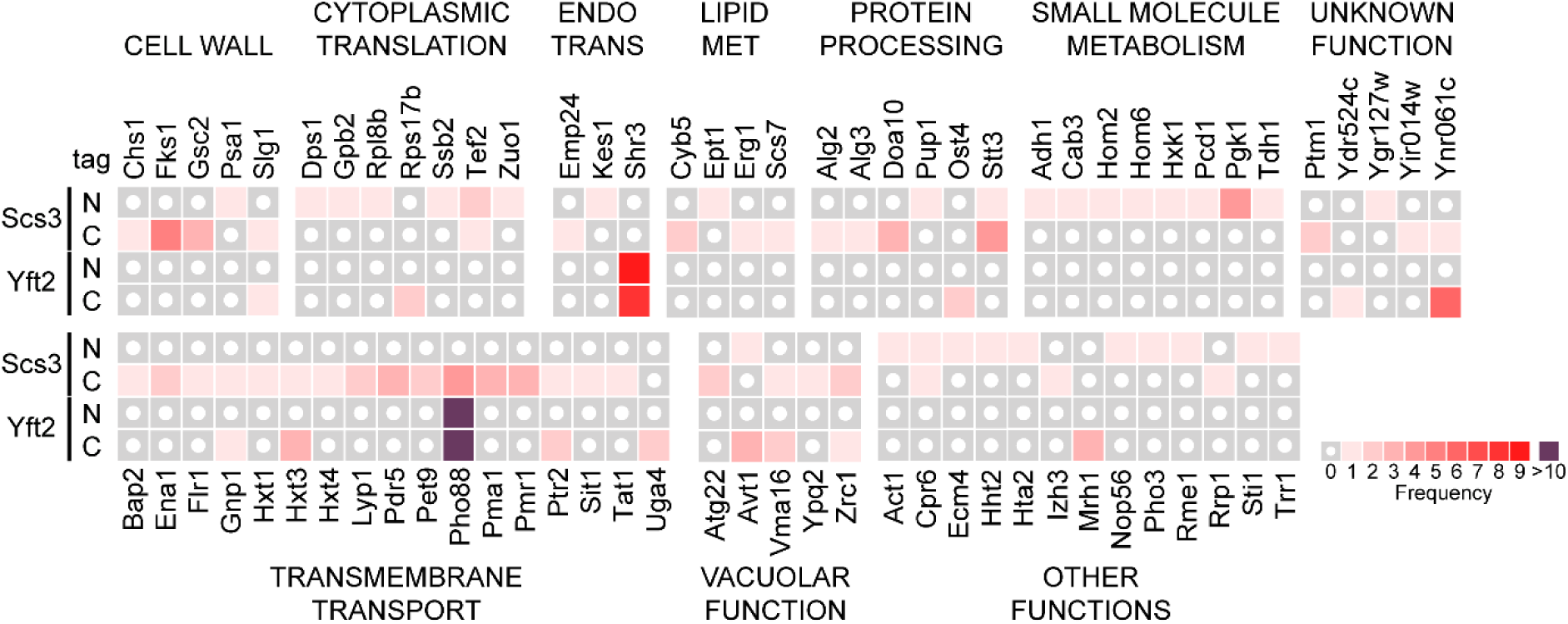
The ScFIT interactome as identified by membrane yeast two-hybrid screening. The frequency by which each unique protein interactor was identified in ScFIT reporter constructs tagged in either N or C-terminus is represented in a heat map and are grouped according to function.

Surprisingly, only few of the ScFIT protein interactors identified with the screen are directly involved in lipid metabolism, suggesting that ScFIT proteins may not function extensively in that cellular process. It could also be noted that while the encoded proteins of genes that *ScFIT* had high degrees of genetic interactions with, such as *ICE2, SEY1* and *UBX2* (14,38), were not identified with the MYTH screen, this does not exclude the presence of a physical interaction. However, this alternatively suggests that they function in parallel but independently in the same cellular process, and that the loss of both is detrimental to cell viability.

Interestingly, several proteins that function in proteostasis and the ubiquitin-proteasome system (UPS) have been found to interact with Scs3 (Fig. 3 and S5). Of note are the J-protein chaperone Zuo1 and the Hsp70 chaperone Ssb2, both of which have been reported to function in protein quality control (39-41). Doa10 is one of the key E3 ubiquitin ligases in yeast, and is involved in ER-associated degradation (ERAD) of proteins (42-44). Lastly, Pup1 is a subunit of the yeast 20S proteasome core (45,46). Taken together, these suggest that Scs3 may function to a certain extent in protein quality control pathways, specifically in the UPS.

### The clearance of ERAD client proteins is impaired in ScFIT mutants

As the accumulation of misfolded proteins and the ensuing proteotoxicity is closely related to the ability of cells to efficiently process client proteins, we hypothesized that the sensitivity to Tm exhibited by *scs3*Δ and *ScFIT*Δ mutants may be the result of impaired protein degradation pathways, as in the case of *ubx2*Δ mutants having impaired turnover of both misfolded ER and cytosolic substrates (47). Following the identification of UPS machinery components as protein interactors of Scs3, we asked if ScFIT plays a role in protein ubiquitination. We overexpressed myc-tagged ubiquitin (Ub-Myc) through and inducible *CUP1* promoter and quantified the extent of total protein ubiquitination in cells by immunoblotting. The amount of ubiquitinated proteins in *ScFIT*Δ mutants was significantly reduced by approximately 38% relative to that of WT cells, with a less pronounced reduction of 18% in *scs3*Δ single mutants (Fig. S6). Since the stabilization of protein substrates was accompanied by a decrease, rather than an increase in high molecular weight ubiquitin antibody-reactive proteins, the inefficient turnover of the said substrates is likely due to a failure to mark them correctly for degradation and not because of efficient clearance in the proteasome.

To investigate if the global decrease of ubiquitinated proteins correlates with protein stability, we measured the turnover of known ERAD substrates in mutants of ScFIT by the cycloheximide chase assay. We expressed HA-tagged Sbh2, Yeh1, and Pgc1 in *ScFIT*Δ mutant. These native proteins are dependent on Doa10-mediated ERAD for normal degradation (42,48). The turnover rates of Sbh2 and Yeh1 in *ScFIT*Δ mutants was similar to WT (Fig. S7A and B). In contrast, the degradation of Pgc1 was significantly accelerated in *ScFIT*Δ mutants (Fig. S7C). Along with the identification of Pgc1 as a Doa10-dependent ERAD substrate, its proper localization dynamics between the ER and LD membranes was found to be critical in determining its stability (48). Doa10 reportedly recognizes ER-localised Pgc1 through its hairpin loop, which then serves as a degron that concentrates Pgc1 on the surface of LDs. As LDs fail to properly mature in the absence of the ScFIT proteins (Fig. 2B), the lateral diffusion of the pool of Pgc1 proteins to the ER may be increased in *ScFIT*Δ mutants, resulting in continual degradation by Doa10 (49). We hypothesized that native proteins in their proper conformation may not illicit a proteotoxic effect on *ScFIT*Δ cells, and that an otherwise compromised protein degradation pathway in this mutant could remain fully capable of clearing these endogenous proteins.

As misfolded model substrate, we monitored the protein levels of epitope-tagged versions of misfolded CPY (CPY*-HA) (Fig. 4A). A small but significant delay in the degradation of CPY* was only observed in *ScFIT*Δ mutants but not in *scs3*Δ nor *yft2*Δ. Next, we measured the degradation rates of the engineered misfolded variant of the Pep4 vacuolar protease (ngPrA*Δ295-331-HA) (50) (Fig. 4B). In contrast to CPY*, *ScFIT*Δ exhibited a strong defect in the degradation of ngPrA*Δ295-331-HA in comparisons to WT and single mutants. Both are luminal soluble substrates which are degraded in a Hrd1-dependent manner (29,50). To further assess if the global decrease of ubiquitinated proteins is associated to ScFIT, we monitored the degradation of San1-dependent cytosolic protein quality control (CytoQC) substrates Δ2GFP-HA and ssPrA-HA (51). Consistent with our results using misfolded ERAD substrates, we found that *ScFIT*Δ mutants are unable to efficiently clear away both cytosolic substrates compared to WT cells or either of the single mutants (Fig. S8). As neither of the single deletion mutants resulted in a stabilization of the ERAD substrates, the two may share a redundant yet poorly understood function. *YFT2* is reported to have been the result of the segmental duplication of *SCS3* (14). This is supported by the more pronounced growth sensitivity to Tm in the double *ScFIT*Δ mutant in comparison to a mild defect in *scs3*Δ cells (Fig. 1C). This, along with the broader range of Scs3 protein interactors (Fig. 3), also suggests of an asymmetric redundancy wherein *YFT2* only partially compensates for the absence of *SCS3* functionality in the ERAD pathway, which ultimately results in less apparent phenotypic defects in *yft2*Δ mutants.

**Figure 4.**
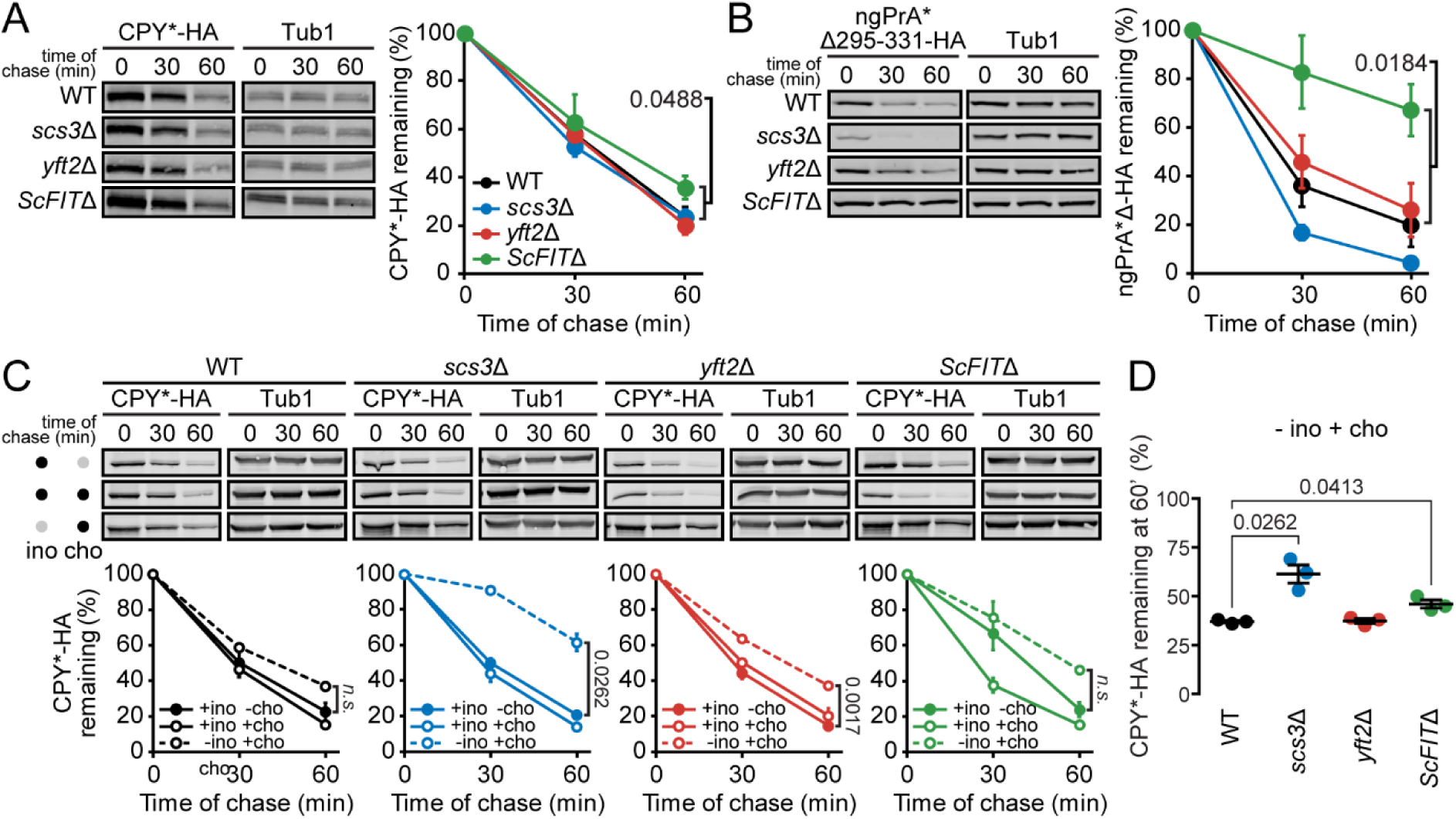
The clearance of ERAD client proteins is impaired in *ScFIT* mutants. (A-B) Protein levels of the model Hrd1-dependent ERAD substrates, (A) CPY*-HA and (B) ngPrA*Δ295-331-HA, were monitored at 0, 30, and 60 min time points following the attenuation of protein translation with cycloheximide. (C) Strains were treated as in (A) but were grown in the presence of inositol and absence of choline (+ ino, - cho), in the presence of inositol and choline (+ ino, + cho), or in the presence of choline and absence of inositol (- ino, + cho). (D) Percentage of CPY*-HA remaining at 60 min of samples grown in the presence of choline and absence of inositol (- ino, + cho). Data shown are the mean ± SEM from three independent experiments. Statistical analysis was subjected to paired two-tailed Student’s t-test.

To further investigate the role of lipid homeostasis and protein quality control, we monitored the degradation of CPY* in *ScFIT*Δ mutant strains supplemented with inositol (+ino, -cho), choline (-ino, +cho), or both (+ino, +cho). There was a significant defect in the degradation of CPY* in *scs3*Δ and *yft2*Δ supplemented with choline compared to inositol (Fig. 4C and D). Similarly, the degradation of CPY* was slower in *scs3*Δ and *ScFIT*Δ compared to WT in the presence of choline. Next, to validate the role of Scs3 in modulating ERAD, we monitored the degradation of ngPrA*Δ295-331-HA in *ScFIT*Δ overexpressing (OE) *SCS3*. In *ScFIT*Δ, ngPrA*Δ295-331-HA was degraded at a significantly slower rate compared to WT cells (Fig. 5A). On the other hand, the degradation of ngPrA*Δ295-331-HA was similar in *ScFIT*Δ OE *SCS3* to WT and *ScFIT*Δ strains, suggesting that *SCS3* is sufficient to rescue the ERAD defect. In the presence of choline (- ino, + cho), the degradation of ngPrA*Δ295-331-HA was decelerated in the three strains (Fig. 5B). Interestingly, OE *SCS3* in *ScFIT*Δ cells supplemented with choline (- ino, + cho) significantly accelerated the degradation of CPY*-HA compared to WT and *ScFIT*Δ (Fig. 5C). As lipid homeostasis correlates with ERAD fitness (7,52), these finding reinforce the notion that ScFIT is essential to regulate lipid levels at the ER and that it contributes to ER proteostasis.

**Figure 5.**
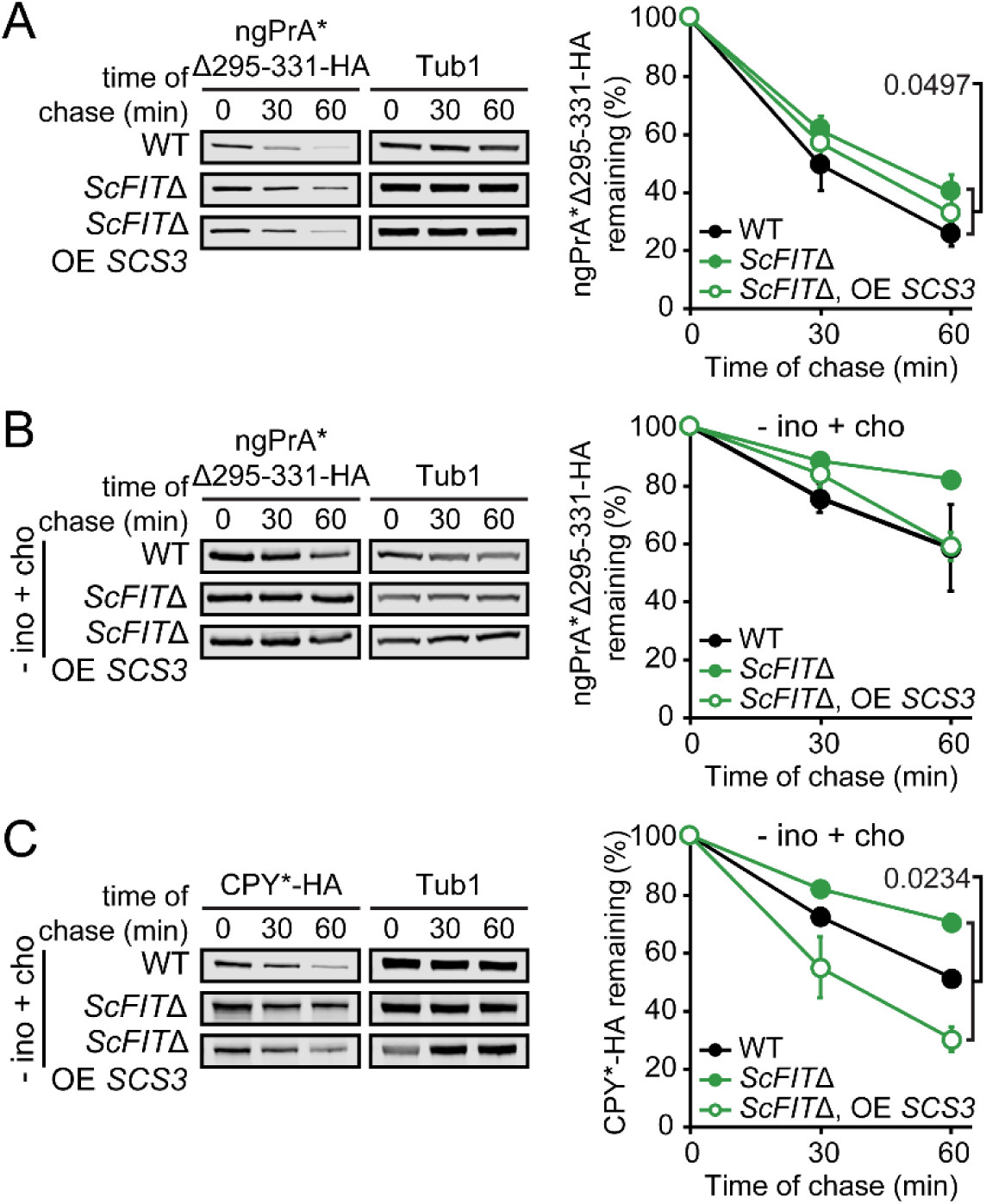
Scs3 is sufficient to rescue proteostatic defect of *ScFIT*Δ. (A) ngPrA*Δ295-331-HA protein levels were monitored at 0, 30, and 60 min time points following the attenuation of protein translation with cycloheximide. OE, overexpressed. (B) ngPrA*Δ295-331-HA protein levels of cells treated as in (A) but in the absence of inositol (- ino, + cho). (C) CPY*-HA protein levels of cells treated as in (A) but in the absence of inositol (- ino, + cho).

## Discussion

Lipid droplets have been increasingly implicated in disease pathophysiology. Despite this, our understanding of its involvement is obscure at best as LD biology is still in its infancy, and more mechanistic insight into LD formation is warranted to grasp its relevance and importance in physiological processes. From the simple budding yeast, several proteins have been identified to influence LD generation (53-55). Among these, the FIT2 class of proteins has similarly gained much interest in recent years, but its initial putative role in LD formation as a lipid-binding protein has recently been contested in favor of a broader function in membrane homeostasis. However seemingly disparate, the identification of lipid phosphatase activity in FIT2 may not be mutually exclusive with previous reports of its involvement in LD biogenesis. Given this, the molecular mechanism by which these two processes are linked is poorly understood, as well as its potential implication for the normal functioning of cells outside the context of LD formation. In this study, we report on the involvement of the yeast FIT homologs (ScFIT) not only in the maintenance of ER membrane homeostasis, but also in coordinating the cellular stress response pathway, namely the UPR, and its consequent impact on protein quality control (Fig. 6).

**Figure 6.**
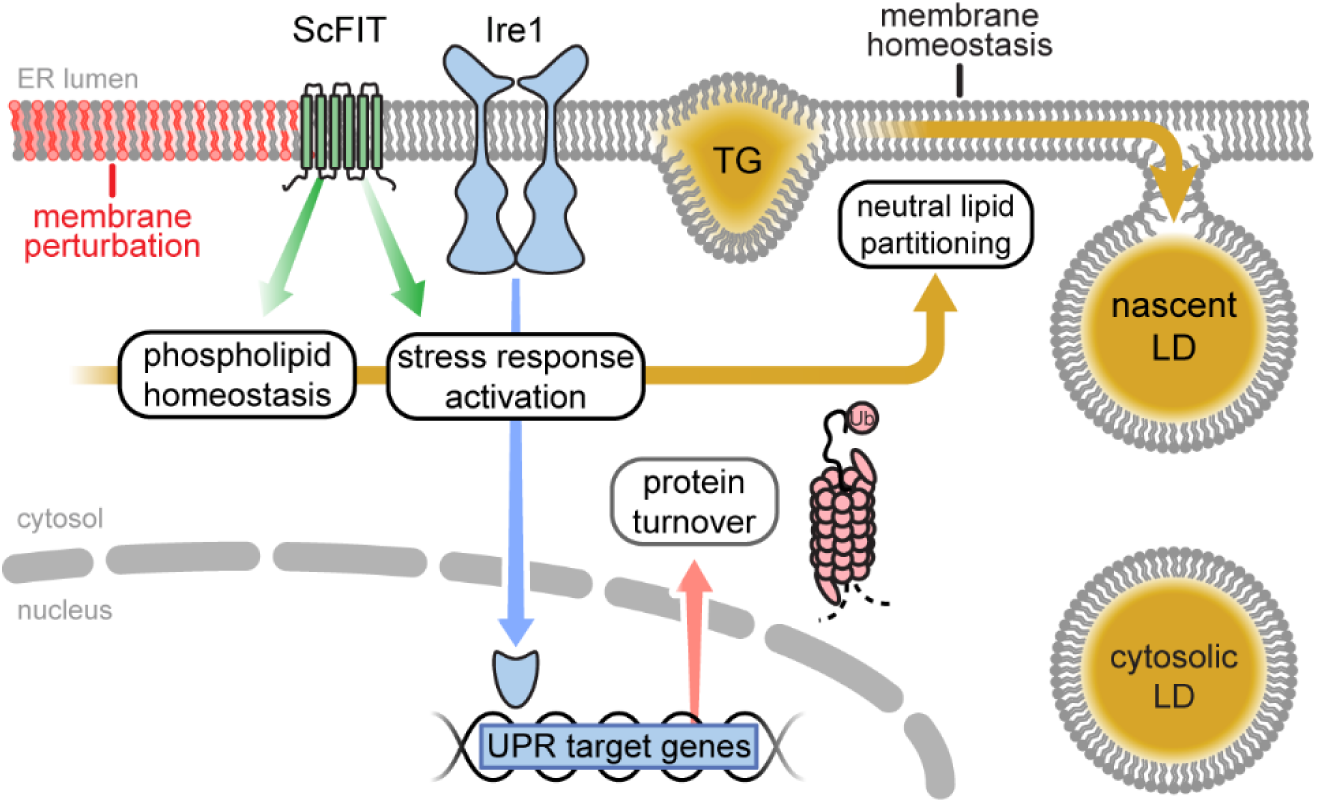
Model for the putative role of ScFIT proteins in coordinating lipid and protein homeostasis. The ScFIT proteins functions in membrane homeostasis by preventing aberrant triacylglycerol (TG) accumulation within the ER, thereby contributing to lipid droplet (LD) maturation. High levels of TG within the ER membrane may impair protein turnover. Alternatively, ScFIT proteins may directly modulate proteostatic pathways, such as the unfolded protein response (UPR) and heat shock response (HSR), to aid in this process with an unknown mechanism.

The complexity of lipid metabolic pathways is underscored by the highly interconnected conversion of intermediates as well as the various organelles and proteins that mediate these processes (56,57). In addition to this, perturbation of the lipid metabolic pathways results in the extensive reprogramming of the bioenergetic network (58,59). Similarly, cellular insults also alter the lipidomic landscape of cells, suggestive of the buffering capacity of lipid pathways against stress conditions. Proteotoxic or lipid bilayer stress (LBS) both activate the UPR and similarly culminate in the formation of LDs. Interestingly, none of the previously reported major protein effectors of LD biogenesis were identified as UPR targets. Moreover, apart from the SE biosynthetic *ARE2*, no other NL synthesis players are upregulated under conditions of ER stress (7,24). Given the recent report of Scs3 as a lipid phosphatase, its consequent upregulation under the UPR program could be part of the effort to orchestrate membrane remodeling.

It was reported that LDs in *ScFIT*Δ cells remain embedded in the ER due to the enrichment of DG, a lipid species with negative membrane curvature, as is the case with the accumulation of PE in a mutant of *CHO2*, a methyltransferase for PC synthesis (20). Interestingly, the addition of either of the positive curvature phospholipids, lyso-PC or lyso-PA, rescued the aberrant LD budding of *ScFIT*Δ cells. The failure of the UPR to restore proper LD maturation in *cho2*Δ cells may result from the markedly reduced capacity to synthesize PC, which exhibits a neutral curvature and is an intermediate to lyso-PC (60). While *cho2*Δ cells indeed accumulated high levels of PE, DG levels are dramatically reduced (7). This taken together with the observation that the gain-of-function DG-binding FIT mutant failed to rescue the aberrant ER membrane whorling in *scs3*Δ cells (19) strongly suggest that defects in ER membrane properties that led to impaired LD maturation may be independent from DG. In contrast, we observed an accumulation of TG at the ER together with irregularly shaped LDs that were detached to the ER in *scs3-1* (Fig. 2B and S4), reinforcing the role of Scs3 in lipid homeostasis.

While the catalytic activity of mammalian FIT2 on PA and lyso-PA was not identified in ScFIT proteins (19,22), this does not exclude the possibility that the latter may instead act on other membrane lipid species *in vivo*. Mammalian FIT2 is proposed to be a lipid phosphate phosphatase enzyme based on *in vitro* evidence (19). As *SCS3* is essential for viability in the absence of a functional UPR (Fig. 1), it can then be hypothesized that Scs3 regulates ER membrane lipid composition, presumably through the conversion of DAG from PA (19), to maintain proper ER function including LD maturation. Taken together, these findings highlight the important role of Scs3 in maintaining ER lipid homeostasis. A unifying model on the role of Scs3 *in vivo* by either acting as an enzyme that catalyzes DAG synthesis, by regulating other enzymes, or by regulating lipid metabolism through other ways should emerge in future studies.

The loss of both FIT homologs in *ScFIT*Δ mutants led to the unexpected stabilization of misfolded proteins in both within the ER and in the cytoplasm (Fig. 4, 5, and S8) which correlate with a global decrease in protein ubiquitination (Fig. S7). The maintenance of ER membrane integrity and lipid homeostasis are critical in supporting organellar function. Loss of *ICE2* exhibits altered ER membrane dynamics including defects in mother-daughter cell ER membrane inheritance and ER-plasma membrane tethering (61,62). This ER membrane perturbation further impaired cellular functions such as phospholipid regulation and protein degradation (38,63-65). Similarly, mutants of the phospholipase Lpl1, which catalyzes the turnover of phospholipids, also exhibited ERAD defects (66,67). The general disruption of lipid metabolism by attenuating FA synthesis in turn caused defects in processing of ERAD client proteins in mammalian systems (68), which may also be in part due to its indirect effects on membrane lipid composition. Conversely, defective protein turnover also exerted a direct effect on membrane composition. The deletion of the ERAD component *UBX2* led to severe changes in ER membrane morphology due to dysregulation of Mga2 processing and the subsequent expression of its transcriptional target *OLE1*, a key regulator of membrane lipid saturation (69). Sterol content within membranes are also under tight control by protein quality control pathways, as the key enzymes Hmg2 and Erg1 are regulated in an ERAD-dependent mechanism (70,71). Intriguingly, these mutants with aberrant membranes exhibited impaired LD formation in addition to curtailed protein turnover (63,67,72).

In a previous study, we have shown that the singular UPR transducer Ire1 in yeast is strongly activated with genetic alterations of ER membrane composition (7). We also identified a LBS sensing switch located at the interface of the amphipathic and transmembrane helices (10) while key residues within the amphipathic helix of Ire1 were reported to be important to sense LBS- and proteotoxic-induced ER stress (27). As ER membrane morphology is compromised in ScFIT mutants, it could be hypothesized that proper Ire1 and UPR activation may not proceed as efficiently, which in turn affects ERAD function. In line with this, several studies lend support for the modulation of LDs by protein quality control pathways including ERAD. The *ubi4*Δ mutant was reported to exhibit less LD accumulation compared to that of WT cells under stress (73). While the rationale for this increase in LD remains enigmatic, it suggests that LD formation may in part be regulated by ubiquitination processes. This agrees with previous studies that detailed on the dependence of the NL biosynthetic enzyme Dga1 and SE lipase Yeh1 on Doa10 for their endogenous turnover (48), and that recruitment of the mammalian ERAD factor UBXD8 onto the LD surface regulates LD growth by modulating lipolysis (74). Apart from LDs, ERAD pathways also regulate the ER membrane composition and phospholipid turnover. The Cdc48 ATPase mediates the processing of the ER membrane sensors Mga2 and Spt24, to yield the cognate transcription factor for *OLE1* regulation (69,75,76), and the degradation of phosphorylation-inactive Pah1 is impaired in proteasome and ubiquitination mutants (77,78). Taken together, these greatly emphasizes the interdependence of membrane homeostasis and protein quality control pathways.

In this study, we build on the current hypothesis on the role of ScFIT proteins in LD formation and membrane homeostasis, and further provide support for its functioning in cell stress response pathways to exert effects on these two processes. (Fig. 6).

## Experimental procedures

### Statistics

Error bars indicate standard error of the mean (SEM), calculated from at least three biological replicates, unless otherwise indicated. *P* values were calculated using one-way ANOVA with Tukey’s post hoc test, unless otherwise indicated and reported as *P* values with 4 significant digits in the figures. All statistical tests were performed using GraphPad Prism 7 software.

### Strains and antibodies

*Saccharomyces cerevisiae* strains used in this study are listed in Table S1. Strains were generated using standard cloning protocols. Anti-HA mouse monoclonal antibody HA.11 (Covance), anti-tubulin mouse monoclonal antibody 12G10 (DHSB), and anti-myc mouse monoclonal antibody (Invitrogen), were commercially purchased. Secondary antibodies goat anti-mouse IgG-DyLight 488 (Thermo Fisher, Waltham, MA), goat anti-mouse IgG-IRDye 800 (LI-COR Biosciences) and goat anti-rabbit IgG-IRDye 680 (LI-COR Biosciences) were commercially purchased.

### Plasmid used in this study

Plasmids and oligonucleotide primers used in this study are detailed in Table S2 and S3, respectively. Plasmids constructs were generated through either conventional restriction enzyme cloning methods or Gibson Assembly (New England Biolabs). The mutant *scs3* library was generated by low-fidelity PCR using primers PS1-PS2 to amplify the promoter, coding sequence, and terminator regions of the *SCS3* from the genomic DNA of wild type (WT) cells. The PCR product was then digested with the enzymes *EcoRI* and *XbaI* before ligation into pGT0004. Plasmid pGT0364 was obtained through a colony sectoring screen detailed in the *Genetic screen for temperature sensitive alleles* section below. Plasmid pGT0286 encoding for WT *SCS3* was similarly generated using conventional PCR amplification. To generate reporter constructs for the membrane yeast two hybrid screen, the coding sequences of *SCS3* and *YFT2* were amplified from WT yeast DNA using primer pairs PS39-PS40 and PS159-PS160, respectively. These were then inserted via Gibson Assembly into vector backbones generated through PCR from pGT0317 using primer pairs PS39-PS40 and PS157-PS158, respectively, to generate pGT0374 and pGT0427. Plasmids pGT0426 and pGT0428 were generated through Gibson Assembly by amplifying the coding sequence of *SCS3* and *YFT2* terminating immediately before the stop codon using WT yeast DNA with primer pairs PS107-PS108 and PS101-PS102, respectively. These were then cloned into PCR-amplified vector backbones using pGT0318 as template with primer pairs PS105-PS106 and PS36-PS99, respectively.

### Spotting growth assay

Strains were grown to saturation in appropriate selective medium overnight at 30°C. Cultures were diluted to 0.2 OD_600_/ml and serially diluted fivefold for a total of four dilutions. The cell suspensions were then spotted onto appropriate agar plates and incubated at indicated temperatures until the appearance of colonies.

### β-galactosidase reporter assay

The β-galactosidase reporter assay was carried out as previously described (29). Typically, cells were grown to early log phase, and tunicamycin (Tm) was added to growth cultures when specified at a concentration of 2.5 µg/ml to cells 1h prior to harvest or a temperature shift to 37°C, for the induction of the UPR and HSR, respectively. Four OD_600_ units of cells were pelleted, washed and resuspended in 75 µl LacZ buffer (125 mM sodium phosphate pH 7, 10 mM KCl, 1 mM MgSO_4_, 50 mM β-mercaptoethanol). An aliquot of 25 μl was transferred into 975 μl ddH_2_O and the absorbance was measured at 600 nm. To the remaining suspension, 50 µl of CHCl_3_ and 20 µl of 0.1% SDS were added and vortexed vigorously for 20 s. The reaction was started with the addition of 700 µl of 2 mg/ml 2-nitrophenyl-β-galactopyranoside (ONPG; Sigma) in LacZ buffer. Next, the reaction was quenched with 500 µl of 1 M Na_2_CO_3_, and total reaction time was recorded. Samples were spun for 1 min at maximum speed. Absorbance of the resulting supernatant was measured at 420 and 550 nm. The β-galactosidase activity was calculated using Eq. (1)

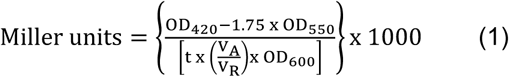

### qPCR

Cells were grown to an early log phase overnight at 30°C. Tunicamycin was added to a final concentration of 2.5 µg/ml and incubated 1h at 30°C or depleted of inositol for 2h, when indicated. Total RNA was extracted using RNeasy Mini Kit (Qiagen) following manufacture’s protocol. DNase treatment in columns was carried out with RNase-free DNase (Qiagen, Venlo, Netherlands) following the manufacturer’s protocol. cDNA was synthesised from 2 μg of total RNA using RevertAid reverse transcriptase (Thermo Fisher, Waltham, MA) following manufacturer’s protocol. SYBR Green qPCR experiments were performed following the manufacturer’s protocol using a QuantStudio 6 Flex Real-time PCR system (Applied Biosystems, Waltham, MA). cDNA (30 ng) and 50 nM of paired-primer mix were used for each reaction. Relative mRNA was determined with the comparative Ct method (ΔΔCt) normalised to housekeeping gene *ACT1*. Oligonucleotide primers used are listed in Table S3.

### Lipid droplets analysis

Cells were grown to early log phase and 500 µL of the suspension was transferred on a coated slide with 10 mg/ml Concanavalin A (Sigma-Aldrich, St. Louis, MO) mounted onto Attofluor cell chamber (Thermo Fisher, Waltham, MA) and imaged at RT. Tunicamycin was added to a final concentration of 2.5 µg/ml and incubated at 30°C for 1h, when indicated. To stain lipid droplets, cells were incubated with 0.05 µg/ml BODIPY 493/503 (Invitrogen) in phosphate buffered saline (pH 7.4) for 10 min at RT, washed and resuspended in liquid medium before transferring into the Attofluor cell chamber for viewing. Samples were imaged with a Leica DMi8 system (HCX PL APO 100x/1.4-0.70 oil immersion objective) under the control of Metamorph Ver. 7.8.10.0, or Zeiss LSM710 Microscope (100× 1.4 NA Plan-Apochromat oil-immersion objective) under the control of Zen (Carl Zeiss MicroImaging).

### Genetic screen for temperature sensitive alleles

The genetic screen was performed as previously reported (30). A mutant library of the *SCS3* open reading frame flanked by 500 bp of its endogenous promoter and 300 bp of its terminator was generated by low-fidelity PCR using Taq DNA polymerase in the presence of 0.05 and 0.1 mM MnCl_2_. DNA fragments were digested with *EcoRI* and *XbaI* and ligate into digested pRS316 to produce a plasmid library *scs3\** with random point mutations. The strain YGT0492 was transformed with the mutant library pRS316-*scs3\** and transformants were spread on selecting synthetic complete lacking uracil (SC-Ura) plates with limiting adenine (low Ade) at 6 µg/ml. Plates were incubated at 25°C until colonies developed fully red pigmentation due to low Ade. Colonies with sectoring phenotype were streaked in duplicate on SC-Ura, low Ade, and incubated at 25°C and 37°C. For the primary screen, 123 colonies were screened for positive clones which sectored at 25°C but remained red at 37°C. From the positive clones, sectoring phenotype at 25°C were re-streaked in duplicate on SC-Ura, low Ade, and incubated at 25°C and 37°C to eliminate false positive. Positive clones were isolated without the plasmid pDN388. Plasmids were extracted from the clones and subjected to DNA sequencing analysis to identify the mutation present in *scs3* temperature sensitive (ts) alleles. The plasmid pGT0364, containing *scs3* ts allele (*scs3-1*) encodes for Scs3 with the following mutated residues D277G and I328V. The plasmid pGT0364 was transformed in strain YGT0492.

### Lipid extraction and fatty acid analysis

Cells were grown to early log phase at 25°C followed by a 2h incubation at 37°C. For whole cell lipid extraction, 10 OD_600_ of cells was washed, pelleted in a glass vial, and lyophilised using Virtis Freezer Dryer under vacuum. All subsequent steps were carried out at 4°C. For lipid extraction of microsomes, 30 OD_600_ of cells was pelleted and resuspended in lysis buffer (50 mM Tris-HCl, pH 7.5, 150 mM NaCl, 5 mM EDTA, 1 mM PMSF and PIC) and lysed mechanically by 15 times of 30 s interval using 0.5 mm zirconium beads at maximum speed of a vortex mixer. The supernatant was collected by spinning down the lysate 5 min at 800 × *g*. The clarified lysate was spun down 1h at 100,000 × *g*. The pellet was resuspended in 100 µl ddH_2_O and sonicated for 30 min before quantifying total protein using the bicinchoninic acid (BCA) protein quantification assay (Sigma-Aldrich). A volume corresponding to 0.3 mg of total protein (79) was transferred into glass vials and lyophilized using Virtis Freeze Dryer under vacuum to record dry weight of each sample. For lipid extraction from whole cells, samples were resuspended in 100 µl ddH_2_O. Afterwards, 300 µl of 0.5 mm zirconium beads and 900 µl of chloroform (CHCl_3_):methanol (2:1) were added before rigorous agitation of 2h at 4°C. From here, 300 µl each of CHCl_3_ and ddH_2_O were added to the mixture and vortexed 15 s twice. The vials were centrifuged 6 min at 4,250 × *g*, and the lower organic phase was transferred to a new glass vial. The extraction step was repeated by the addition of 500 µl of CHCl_3_ and further agitation 2h. Lipid extraction from microsomes was done similarly with scaled-down reagent volumes. Combined extracts were concentrated, resuspended in 50 µl (20 µl for microsomal extracts) CHCl_3_:methanol (2:1), and spotted on HPTLC Silica gel 60 plates (Merck Millipore) using Linomat 5 (CAMAG). Triacylglycerol (TG) were separated with toluene:CHCl_3_:methanol (85:15:5) and visualized under long-wave ultraviolet light (320 nm) by spraying 0.05 mg/ml of primuline dye in acetone:water (80:20) onto the dried plates.

Spots corresponding to TG were scraped off the silica plates and transferred into glass vials. A total of 100 µl 1 mM pentadecanoic acid (C15:0) was added to the tubes as internal standard. TGs were hydrolysed and esterified to fatty acid methyl esters (FAME) with 300 µl of 1.25 M HCl-methanol for 1h at 80°C. FAMEs were extracted three times with 1 ml of hexane. Combined extracts were dried under nitrogen, resuspended in 20 µl. FAMEs were separated by gas chromatography with a flame ionisation detector (GC-FID; GC-2014, Shimadzu, Kyoto, Japan) using an ULBON HR-SS-10 50 m × 0.25 mm column (Shinwa, Tokyo, Japan). Supelco 37 component FAME mix was used to identify corresponding FAs (Sigma-Aldrich, St Louis, MO). Data were normalised using the internal standard C15:0.

### Transmission Electron Microscopy

Samples for transmission electron microscopy (TEM) were prepared as previously described (80). One OD_600_ unit of early log-phase cells grown at 25°C or 37°C was collected and pre-fixed with glutaraldehyde overnight at 4°C. Post-fixation was performed in the presence of 2% potassium permanganate for 1h at room temperature. After dehydration in ethanol, cells were infiltrated with Spurr’s resin and incubated for 24h at 60°C to allow polymerization. Silver-gray sections were prepared using Ultracut UCT (Leica) equipped with a diamond knife and stained with lead citrate. Micrographs were taken using a transmission electron microscope (Joel JEM-1230).

### Phospholipid analysis

Cells were grown to mid-log phase overnight in medium containing 75 µM inositol with (+cho) or without (-cho) 1 mM choline in the presence of 10 µCi/ml [^32^P]-orthophosphate. Cells collected by filtration were resuspended in medium with or without inositol or choline, as indicated, and in the presence of 10 µCi/ml [^32^P]- orthophosphate. Cells were harvested after 3h following the shift or at indicated time point. Labelled lipids were extracted as previously described (81). The individual phospholipid species were resolved by two-dimensional thin layer chromatography. Phospholipids were separated using the solvent system chloroform/ ethanol/water/triethylamine (30:35:7:35) for at least 2h. Phospholipid identity was based on the mobility of known standards and quantified on a STORM 860 PhosphorImager (Amersham Biosciences).

### Cycloheximide chase assay

The cycloheximide chase assay was performed as previously described (51). In brief, 6 OD_600_ units of early log phase cells were grown in appropriate selective media. To induce lipid perturbation, 1 mM of choline chloride was added a day prior to harvesting, whereas inositol depletion was performed two hours beforehand. Protein synthesis was inhibited by adding 200 μg/ml cycloheximide. Samples were taken at designated time points and trichloroacetic acid (TCA) was added to a final 10% volume. Cells were mechanically disrupted with 300 µl of 0.5 mm zirconium beads at 6,500 rpm for 2 × 30 s using a tissue homogeniser (Precellys 24, Bertin Instruments). Precipitated proteins were pelleted 10 min at 21,000 × *g*, 4°C, and resuspended in 40 µl of TCA Resuspension Buffer (100 mM Tris pH 11, 3% SDS, 1 mM PMSF, PIC). Solubilised proteins were separated by SDS-PAGE, transferred on nitrocellulose membranes. Immunoblotting was performed with appropriate primary antibodies and IRDye-conjugated secondary antibodies. Proteins were visualised using the NIR fluorescence system (Odyssey CLx Imaging System). Values for each time point were normalised using anti-Tub1 as loading controls. Tonal quality was adjusted for representative images through ImageStudio Lite Version 5.2 (LI-COR Biosciences) where appropriate and was followed by quantification. All comparative analyses were done on immunoblots performed in parallel using samples derived from the same experiment.

## Acknowledgements

We are grateful to Dr. Davis Ng for providing reagents. We thank to members of Thibault lab for critical reading of the manuscript.

## Conflict of interest

The authors declare that they have no conflicts of interest with the contents of this article.

## Author contributions

Conceptualisation: G.T; Methodology: P.J.S., W.S.Y., M.L.G., C.M., S.A.H. and G.T.; Formal Analysis: P.J.S., W.S.Y., and M.L.G.; Investigation: P.J.S., W.S.Y., M.L.G., S.A.J., and C.M.; Resources: P.J.S., W.S.Y., and M.L.G.; Writing – Original Draft: P.J.S. and G.T.; Writing – Review & Editing: P.J.S., W.A.P., M.L.G., W.A.P., and G.T.; Funding Acquisition: P.J.S., W.A.P., S.A.H., and G.T.; Supervision: G.T.

## FOOTNOTES

This work was supported by the Nanyang Assistant Professorship programme from the Nanyang Technological University to G.T., the National Research Foundation, Singapore, under its NRF-NSFC joint research grant call (Synthetic Biology, NRF2018NRFNSFC003SB-006) to G.T., the Nanyang Technological University Research Scholarship to P.J.S. (predoctoral fellowship), the NIH grant GM-19629 to S.A.H. the Intramural Research Program of the NIH to W.A.P., The National Institute of Diabetes and Digestive and Kidney Diseases (NIDDK) to W.A.P.

## The abbreviations used are

3’-AT: 3’-amino-1,2,4,-triazole
CD4: cluster of differentiation 4
CytoQC: cytosolic protein quality control
DAG: diacylglycerol
ER: endoplasmic reticulum
ERAD: ER-associated degradation
FIT: fat storage inducing transmembrane
GC-FID: gas chromatography with flame ionization detector
HSR: heat shock response
LBS: lipid bilayer stress
LD: lipid droplet
MAG: monoacylglycerol
MYTH: membrane yeast two hybrid
PA: phosphatidic acid
PC: phosphatidylcholine
PE: phosphatidylethanolamine
PI: phosphatidylinositol
PS: phosphatidylserine
OE: overexpression
ScFIT: *S. cerevisiae* FIT homologues
SE: steryl ester
SGA: synthetic genetic array
TEM: transmission electron microscopy
TG: triacylglycerol
TLC: thin layer chromatography
Tm: tunicamycin
UPR: unfolded protein response
UPS: ubiquitin-proteasome system
X-gal: 5-bromo-4-chloro-3-indolyl-β-D-galactopyranoside.

